# Restrictive eating is linked to weaker recollection of food in young adults

**DOI:** 10.64898/2026.11.12.766220

**Authors:** Chong Zhao, Jennifer E. Wildes, Akram Bakkour

**Affiliations:** Department of Psychology, University of Chicago, Chicago, Illinois 60637, USA; Institute for Mind and Biology, University of Chicago, Chicago, Illinois 60637, USA; Department of Psychiatry and Behavioral Neuroscience, University of Chicago, Chicago, IL 60637, USA; Neuroscience Institute, University of Chicago, Chicago, Illinois 60637, USA

## Abstract

Affective significance often enhances episodic memory, with stronger effects on recollection than on familiarity, but prior research has focused primarily on within-person manipulations of emotional valence. Here, we tested a novel question: whether stable individual differences in the personal significance of food were associated with long-term memory across individuals, given that food ranges from highly appetitive to strongly avoided. Across two experiments, we examined familiarity-based memory (using an image recognition task) and recollection-based memory (using a spatial source recognition task) for food and nonfood stimuli as a function of eating-related individual differences in young adults. In Experiment 1, familiarity-based recognition memory was largely unrelated to eating behavior. In Experiment 2, however, greater restrictive eating and fear of negative consequences from eating were associated with poorer recollection-based memory for food when compared to nonfood stimuli. Together, these findings suggest that individual differences in the affective significance of food do not correlate with episodic memory uniformly but instead are selectively associated with reduced recollection, but not familiarity, of food relative to neutral stimuli.

## Introduction

Episodic long-term memory refers to memory for personally experienced events, including the contextual details that specify what happened, where it occurred, and when it took place. Dual-process models propose that episodic retrieval is supported by two distinct processes: recollection and familiarity (Yonelinas, 2001). Recollection involves the retrieval of contextual details associated with a prior event, whereas familiarity reflects a context-free sense that an item has been encountered before. Although familiarity supports recognition memory, tasks that require memory for contextual features, such as source memory paradigms, depend primarily on recollection (Yonelinas, 1999).

This distinction is important because episodic memory performance may be shaped by differences in affective salience. Among the range of episodic memories people store, positively and negatively valenced memories are often more accessible than neutral memories (Williams et al., 2022). Consistent with this, a substantial body of research has shown that emotionally salient information is remembered better than neutral information. Importantly, this emotional memory advantage appears to be driven largely by recollection rather than familiarity, suggesting that affective significance particularly enhances the retrieval of contextual details from past events. For example, Ochsner (2000) reported enhanced recollection, but not familiarity, for negative relative to neutral stimuli. Converging neural evidence further supports this account: negative faces elicit a larger parietal old/new ERP effect, a well-established neural correlate of recollection (Johansson et al., 2004; Rugg & Curran, 2007).

Importantly, this literature has almost exclusively manipulated emotional valence within individuals, comparing memory for positive, negative, and neutral stimuli presented to the same participants. Consequently, it remains unclear whether the memory advantage associated with affective significance also extends to stable between-person differences. Specifically, if a category of stimuli carries greater personal affective significance for some individuals than others, does this translate into enhanced long-term memory across individuals?

We chose food as our stimulus category because it combines broad relevance with substantial individual variability in personal meaning. Food is a universally important class of stimuli, yet people’s relationships with food differ markedly, ranging from strong preferences and enjoyment to concerns about dieting, disgust, or restrictive eating (i.e., severely limiting the type and/or quantity of food consumed). This variability makes food especially well-suited for examining how affective significance shapes memory across individuals. Consistent with this idea, recent research has increasingly used food as a domain for studying visual and cognitive performance, showing that food-related individual differences, such as food neophobia and disgust, are meaningfully linked to recognition and perceptual abilities (Gauthier & Fiestan, 2023; Gauthier & Olatunji, 2024; Smithson et al., 2026). In the present study, individual differences in eating-related experience and attitudes were assessed using eating behavior questionnaires (Burton Murray et al., 2021; Forbush et al., 2013), allowing us to test whether the effect of affective significance on memory varies across people.

To clarify the possible ways affective significance may influence memory, we propose a conceptual triad of hypotheses. First, the **Memory Facilitation Hypothesis**, suggests that personally significant stimuli naturally attract greater mnemonic engagement at encoding, resulting in enhanced episodic recollection. For instance, events with positive emotions led to more integrated episodic memories in lab and real world settings (Pratt et al., 2025). This perspective argues that memory benefits are not just a product of manipulated emotional states in the lab but extend to enduring individual differences. Thus, individuals for whom food is more emotionally salient, whether due to strong preferences or heightened concerns, should exhibit superior memory for food-related events and details.

Conversely, the **Memory Independence Hypothesis** posits that the affective dimensions of eating behavior are largely dissociated from episodic memory. For example, when participants were asked to study positive, neutral, and negative picture stimuli, their memory performance did not differ as a function of affect when they deeply encoded the images through a writing task (Leventon et al., 2018). Although people differ markedly in how much they care about food and in their approach to eating, these affective distinctions do not necessarily translate to differences in encoding or retrieval. According to this model, the recollection advantage observed for emotional stimuli in laboratory studies is a consequence of transient states, not lasting personal relevance; individual differences in eating behavior are therefore expected to leave memory performance unaffected.

Finally, the **Memory Suppression Hypothesis** introduces the possibility that personally salient food stimuli may elicit cognitive suppression or avoidance, especially among those with restrictive eating tendencies. For instance, negative affect has been shown to impair memory for contextual information in a paired-associate task (Bisby & Burgess, 2014). Thus, it is plausible that rather than enhancing food-related memory, for individuals with restrictive eating affective significance instead motivates regulatory strategies that reduce the encoding or retrieval of food-related information. As a result, their heightened conflict or concern surrounding food may lead to poorer episodic memory, contrasting with the memory-enhancing effects typically associated with emotional salience.

By situating food at the intersection of personal relevance and emotional significance, this framework allows us to examine whether the mnemonic effects of affect extend beyond laboratory manipulations to the stable, meaningful experiences of everyday life. Through these hypotheses, we aim to characterize not just if, but how, affective significance, shaped by eating behavior, modulates episodic memory.

### The Present Study

Our study was designed to investigate whether individual differences in affective preferences for food, as measured by indices of restrictive eating, influence performance on two different components of episodic long-term memory: familiarity in Experiment 1 and recollection in Experiment 2. Specifically, we evaluated three competing hypotheses, the **Memory Facilitation Hypothesis**, the **Memory Independence Hypothesis**, and the **Memory Suppression Hypothesis**. Across both experiments, if individuals with more restrictive eating tendencies demonstrated selectively improved memory for food items compared to non-food items, this was interpreted as support for the **Memory Facilitation Hypothesis**. Conversely, if no relationship emerged between eating behavior and memory performance, this was viewed as being consistent with the **Memory Independence Hypothesis**. Lastly, if restrictive eating was associated with poorer memory for food stimuli, this pattern favored the **Memory Suppression Hypothesis**.

Experiment 1 focused on familiarity-based memory. Participants first completed a working memory task involving both food and non-food stimuli, followed by a recognition task intended to target the familiarity component of long-term memory. Recognition memory was operationalized as participants’ ability to correctly identify previously encountered items, primarily relying on familiarity rather than contextual detail. By comparing performance across these tasks, we evaluated whether eating preference interacts with stimulus category in a manner consistent with the hypothesized models.

Experiment 2 addressed recollection-based memory. In this experiment, participants studied images of food and non-food items, each paired with a unique spatial location. On subsequent tests, participants were asked to recall the spatial location associated with each item, a process that requires the retrieval of context and thus indexes recollection more directly. To assess whether learning trajectories differed as a function of eating behavior, encoding and retrieval procedures were repeated across five learning-test cycles, and spatial recall was evaluated on the following day. This design permitted a thorough investigation of whether restrictive eating behaviors modulate recollection for food stimuli and shape the slope of learning over time.

Altogether, Experiment 1 tested whether restrictive eating behavior affects familiarity-based recognition memory for food stimuli compared to nonfood stimuli. Experiment 2 extended this approach to recollection, measuring spatial source memory and learning curves for food compared to nonfood in the context of eating preferences.

## Experiment 1

### Methods

#### Participants

Participants were recruited via Prolific using the following inclusion criteria: ages 18–35, current residence in the United States, fluency in English, normal or corrected-to-normal vision, no colorblindness, and no self-reported psychological or neurological disorders. To ensure participant engagement and data quality, five attention check questions were embedded throughout the online task. These consisted of straightforward classification items (e.g., “Is a coffee mug edible?”), with participants instructed to press ‘E’ for edible or ‘I’ for inedible. Attention check items have been validated in previous studies employing the change localization task (Zhao et al., 2022). Only participants who answered all five attention checks correctly were included in the final analyses. A sample size estimation was conducted using G*Power, based on a moderately strong expected effect size (Cohen’s d = 0.7), a significance level of .05, and 80% power. Results indicated that a minimum of 68 participants would be sufficient to detect group differences reliably. Accordingly, we recruited 89 young adults (ages 18-35, M = 28.97, SD = 4.27) meeting all eligibility criteria to ensure adequate power for hypothesis testing. Gender identity was distributed as Female (53.93%), Male (44.94%), and Other (1.12%). Racial identity was distributed as White/Caucasian (64.04%), Black or African American (20.22%), Asian (10.11%), More than one race/ethnicity (3.37%), American Indian or Alaskan Native (1.12%), and Choose not to respond (1.12%). Study procedures were approved by the relevant University of Chicago Institutional Review Board and participants were compensated for their participation.

#### Stimuli

Food stimuli were selected from a validated set of food images from the Food-Pics database (Blechert et al., 2019), which provides standardized images of diverse food items. Control stimuli consisted of images of animals, sourced from the object set used in Brady et al. (2008). All images were presented against a uniform gray background (RGB: 128, 128, 128) with a central black fixation cross (30 px, Arial font).

For the working memory task, each trial displayed images on a 1000 × 800 pixel canvas. Stimulus images, either food or neutral (animal), were rendered at 200 × 200 pixels and presented at one of four quadrant locations on the canvas. The location assignments were randomized across trials, and participants were instructed to maintain fixation on the central cross throughout.

During the long-term memory (LTM) task, images (200 × 200 pixels) were centered on the same gray canvas background. The fixation cross was displayed until the presentation of the stimulus. Participants responded to memory and affective rating prompts while the stimulus was shown centrally. All visual elements were presented using the jsPsych canvas-keyboard interface.

#### Procedure

Participants completed two tasks in sequence: a visual working memory task followed by a long-term memory retrieval and affective rating task. The working memory task utilized a change detection paradigm with set sizes of 2 or 4 items, using both food and neutral stimuli (Fukuda & Vogel, 2019). On each trial, a fixation cross appeared for 250 milliseconds, followed by the stimulus display for 500 milliseconds. Participants were then given a 1,000-millisecond retention interval to maintain the information in working memory. After the delay, the probe display appeared, and participants indicated whether the stimulus had changed or remained the same. Intertrial intervals were set at 500 milliseconds, and responses were collected until the participant made their choice. Each participant completed 120 trials, organized across set sizes and stimulus categories.

Following the working memory phase, participants completed a long-term memory retrieval task. In this portion, each trial began with a fixation cross and was followed by the presentation of a single image. Participants indicated whether they had previously seen the image during the working memory task (old/new decision) and were allowed unlimited time to respond. This task consisted of 120 old (previously seen) and 120 new images. Immediately after the memory judgment, participants rated each image on an affective scale ranging from “pleasant” to “unpleasant.” The affective rating responses were also collected until the participant responded.

#### Restrictive Eating Measures

Individual differences in the affective salience of food were assessed using the Eating Pathology Symptoms Inventory (EPSI; Forbush et al., 2013) and the Nine-Item ARFID Screen (NIAS; Burton Murray et al., 2021). The EPSI is a multidimensional self-report measure designed to assess a broad range of eating-disorder symptoms across diverse populations. It comprises eight subscales, with higher scores indicating more severe eating pathology. In the current study, we used two EPSI subscales that reflect different facets of restrictive eating. The Restricting subscale comprises six items that measure behavioral facets of dietary restriction, particularly the extent to which individuals limit their food intake in relation to weight and shape concerns. In contrast, the 3-item Cognitive Restraint subscale assesses the degree to which individuals intend to limit or control their food intake, whether or not these attempts are successful. The EPSI has demonstrated excellent internal consistency and test-retest reliability in previous research (Forbush et al., 2013). Both EPSI subscales demonstrated acceptable to good internal consistency (Restrictive Eating: Cronbach’s alpha = 0.857; Cognitive Restraint: Cronbach’s alpha = 0.745).

The NIAS is a self-report measure used to screen for symptoms of avoidant/restrictive food intake disorder (ARFID). It is comprised of three subscales (Picky Eating, Appetite, and Fear) and a total score that reflects overall ARFID symptom severity. The Picky Eating subscale measures sensory aversion to food; the Appetite subscale measures a lack of interest in eating or food; and the Fear subscale measures fear of aversive consequences of eating such as choking or vomiting. Unlike the EPSI Restricting and Cognitive Restraint subscales, ARFID-related food restriction is not primarily motivated by weight or shape concerns. Thus, the NIAS captures a conceptually distinct form of restrictive eating from the EPSI subscales. NIAS subscale scores range from 0-15, and the total score ranges from 0 to 45, with higher scores indicating more severe ARFID symptoms. In addition, validated clinical cutoffs of 10 for the Picky Eating and Fear subscales and 9 for the Appetite subscale have been reported (Burton Murray et al., 2021). The NIAS total score showed strong internal consistency across the board, with high reliability for the total scale (Cronbach’s alpha = 0.862) and each of its subscales (Picky Eating: alpha = 0.835; Appetite: alpha = 0.842; Fear of Negative Consequences: alpha = 0.923).

**Table 1.**
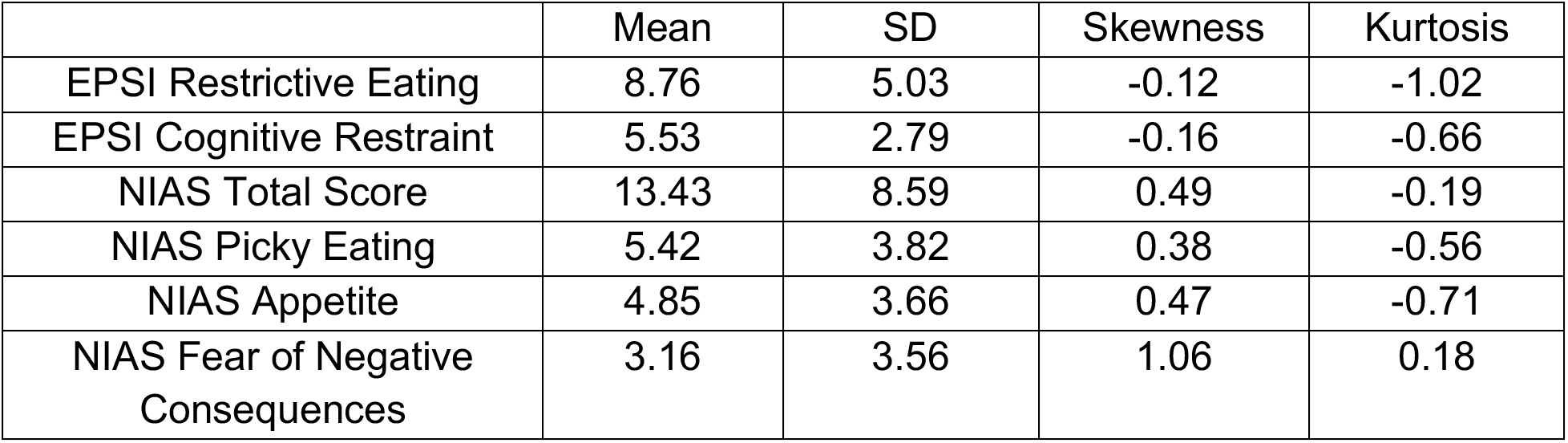
Mean, SD, Skewness and Kurtosis of Eating-Related Symptoms in Exp 1.

|  | Mean | SD | Skewness | Kurtosis |
|---|---|---|---|---|
| EPSI Restrictive Eating | 8.76 | 5.03 | -0.12 | -1.02 |
| EPSI Cognitive Restraint | 5.53 | 2.79 | -0.16 | -0.66 |
| NIAS Total Score | 13.43 | 8.59 | 0.49 | -0.19 |
| NIAS Picky Eating | 5.42 | 3.82 | 0.38 | -0.56 |
| NIAS Appetite | 4.85 | 3.66 | 0.47 | -0.71 |
| NIAS Fear of Negative Consequences | 3.16 | 3.56 | 1.06 | 0.18 |

### Results

#### Restrictive eating was associated with reduced affective ratings for food relative to nonfood stimuli

To examine whether eating-related symptoms influenced affective ratings for food relative to nonfood stimuli, we first tested whether individual differences in eating behavior were associated with differences in affective responses across stimulus categories. We conducted a series of mixed-effects linear models predicting affective ratings from stimulus category (food vs. nonfood), eating-related symptoms, and their interaction.

Across all three eating-related measures, EPSI Restricting, EPSI Cognitive Restraint, and NIAS total score symptoms, there was a consistent main effect of stimulus category, such that nonfood stimuli were rated less positively than food stimuli. This effect was significant in the EPSI Restricting model (b = -0.418, SE = 0.064, z = -6.56, p < .001, 95% CI [-0.543, -0.293]), the EPSI Cognitive Restraint model (b = - 0.314, SE = 0.072, z = -4.39, p < .001, 95% CI [-0.454, -0.173]), and the NIAS total score model (b = -0.421, SE = 0.059, z = -7.16, p < .001, 95% CI [-0.536, -0.306]).

Critically, the results most relevant to our prediction were the significant Category × Restricting and Category × NIAS total score interactions, which indicate that eating-related symptoms were specifically associated with altered affective responses to food relative to nonfood stimuli. EPSI Restricting significantly predicted lower affective ratings overall (b = -0.009, SE = 0.004, z = -2.12, p = .034, 95% CI [-0.018, -0.001]), but more importantly, this effect was qualified by a significant Category × Restricting interaction (b = 0.022, SE = 0.006, z = 3.41, p = .001, 95% CI [0.009, 0.034], Fig. 2), indicating that the negative association between restrictive eating and affective ratings was stronger for food than for nonfood stimuli. A similar pattern emerged for ARFID symptoms. Although ARFID symptoms were associated with lower affective ratings overall (b = -0.007, SE = 0.003, z = -2.77, p = .006, 95% CI [-0.012, -0.002]), the key finding was the significant Category × NIAS total score interaction (b = 0.014, SE = 0.004, z = 3.86, p < .001, 95% CI [0.007, 0.021], Fig. 2), again showing that higher ARFID symptoms were more strongly associated with reduced affective ratings for food than for nonfood stimuli. These interaction effects are central to our hypothesis because they suggest that restrictive eating is not simply linked to lower affective ratings in general, but rather to a food-specific reduction in affective response. In contrast, EPSI Cognitive Restraint neither significantly predicted affective ratings overall (b = -0.011, SE = 0.008, z = -1.33, p = .182, 95% CI [-0.027, 0.005]) nor interacted significantly with category (b = 0.015, SE = 0.012, z = 1.32, p = .187, 95% CI [-0.007, 0.038], Fig. 2), providing no evidence that it was associated with differential affective responses to food versus nonfood stimuli. Taken together, these findings indicate that food stimuli were rated more positively than nonfood stimuli and that higher levels of EPSI Restricting and ARFID symptoms were associated with lower affective ratings, particularly for food cues.

**Fig 2.**
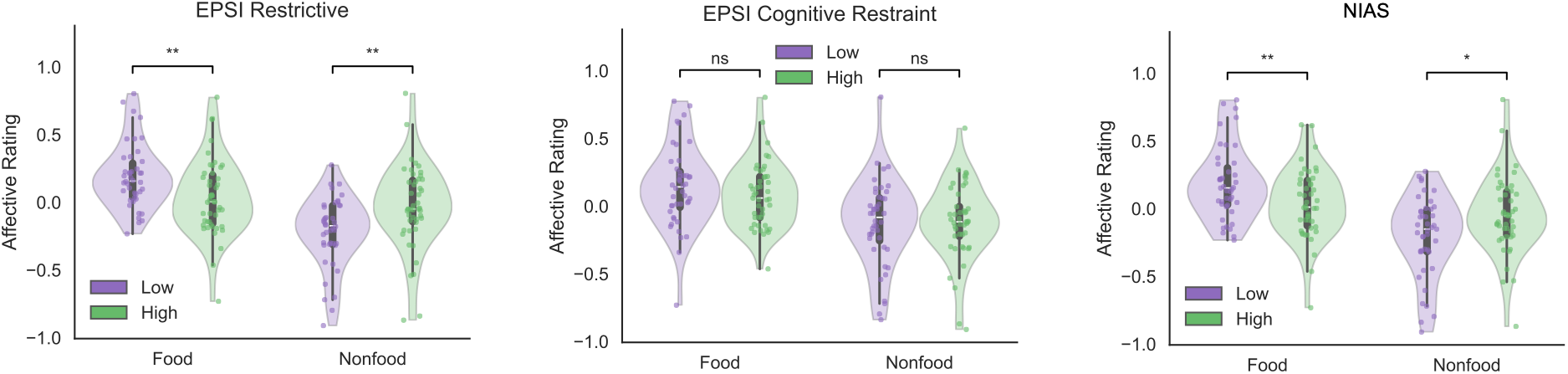
Restrictive eating behavior reduces affective ratings for food relative to nonfood stimuli. Participants with high scores on EPSI Restricting showed lower affective ratings for food than for non-food images (left panel), whereas there was no difference in affective ratings between food and non-food images for either the high or low EPSI Cognitive Restraint groups (middle panel). Participants with high NIAS scores showed selectively reduced affective ratings for food relative to non-food images (right panel).

#### Restrictive eating and Cognitive Restraint were not correlated with recognition memory for food relative to nonfood stimuli

Given that eating-related symptoms were associated with food-specific differences in affective ratings, we next turned to our primary question of whether these differences would be reflected in recognition memory, a measure that primarily taps familiarity. Within the proposed framework, support for the **Memory Facilitation Hypothesis** would require evidence that greater restrictive eating tendencies were associated with better recognition memory for food than for nonfood items. Support for the **Memory Suppression Hypothesis** would require the opposite pattern, such that greater restrictive eating tendencies predicted poorer recognition memory specifically for food stimuli. By contrast, the **Memory Independence Hypothesis** would be supported if eating-related traits were unrelated to recognition memory performance and did not interact with stimulus category.

**Fig 3.**
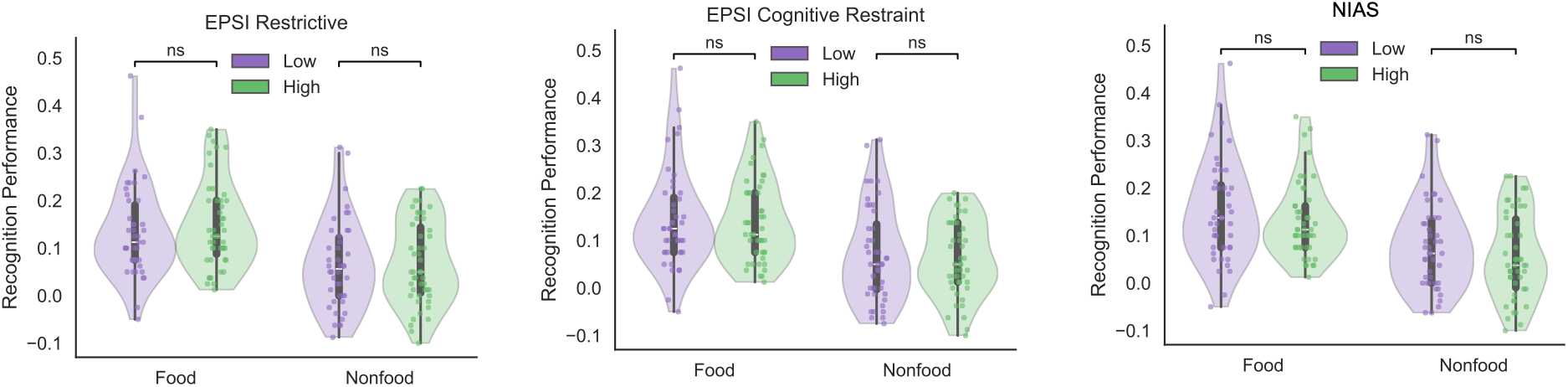
Restrictive eating behavior did not affect recognition memory performance for food relative to nonfood stimuli. There were no differences in recognition memory performance for food versus non-food images between the high and low groups on EPSI Restricting, EPSI Cognitive Restraint, or ARFID symptom severity (measured with NIAS score).

The results were most consistent with the **Memory Independence Hypothesis**. Across the mixed linear models, recognition memory performance, indexed as hits minus false alarms, was not significantly associated with EPSI Restricting (b = -0.000, SE = 0.004, z = -0.10, p = .922), and Restricting did not interact with stimulus category (b = 0.000, SE = 0.006, z = 0.06, p = .952) or with encoding set size and category in a three-way interaction (b = -0.000, SE = 0.002, z = -0.27, p = .789). EPSI Cognitive Restraint also did not significantly predict recognition memory (b = -0.009, SE = 0.008, z = -1.24, p = .216), and its interaction with category was not significant (b = 0.017, SE = 0.010, z = 1.66, p = .098). Similarly, ARFID symptoms were not significantly related to recognition memory performance (b = -0.003, SE = 0.002, z = -1.04, p = .297), nor did they interact with category (b = 0.002, SE = 0.003, z = 0.70, p = .487) or with category and set size jointly (b = -0.001, SE = 0.001, z = -0.97, p = .333). To further examine whether individual differences in ARFID symptoms moderated the effect of category on performance, we conducted separate mixed-effects models for the three ARFID subscales: Fear, Picky Eating, and Appetite. Across all three models, there was no evidence that the association between category (food vs. nonfood) and performance varied as a function of ARFID symptoms. Specifically, the Category × NIAS Fear interaction was not significant (b = 0.026, SE = 0.024, z = 1.08, p = .279). Similarly, neither the Category × NIAS Picky Eating interaction (b = 0.006, SE = 0.023, z = 0.27, p = .788), nor the Category × NIAS Appetite interaction (b = 0.007, SE = 0.024, z = 0.30, p = .765) was significant. Furthermore, all of the three-way interactions between Category, Set Size, and each ARFID subscale were not significant (NIAS Fear: b = −0.010, SE = 0.008, z = −1.36, p = .174; NIAS Picky Eating: b = −0.004, SE = 0.007, z = −0.59, p = .558; NIAS Appetite: b = −0.003, SE = 0.008, z = −0.34, p = .732). Thus, none of the eating-related predictors showed the critical selective association with food memory that would support either the Memory Enhancement or Memory Suppression account.

The only reliable effect to emerge was a main effect of stimulus category in the Cognitive Restraint model, indicating that recognition memory performance was lower for nonfood than for food stimuli (b = -0.146, SE = 0.064, z = -2.28, p = .023). However, because this effect was not qualified by Cognitive Restraint or any other eating-related individual difference, it does not provide evidence for either enhancement or suppression as a function of eating behavior. In addition, encoding working memory set size did not significantly predict recognition performance in any model (ps ≥ .801), nor did it interact with category or eating-related symptoms. Taken together, these findings suggest that, in Experiment 1, familiarity-based recognition memory was largely independent of individual differences in restrictive eating and related eating symptom measures, providing the strongest support for the **Memory Independence Hypothesis**. However, because Experiment 1 tested primarily familiarity-based recognition rather than recollection, it remained possible that eating-related differences would emerge for a stronger, more detail-dependent form of memory. We therefore conducted Experiment 2 using a source memory task to test recollection more directly.

## Experiment 2

### Methods

#### Participants

Participants were recruited via Prolific with the same criteria as Experiment 1. We enrolled 70 young adults (ages 18-35, M = 29.22, SD = 4.50) meeting all eligibility criteria to ensure adequate power for hypothesis testing. The gender distribution was 61.43% Female (n = 43), 35.71% Male (n = 25), and 2.86% Other (n = 2). Regarding racial identity, participants identified as White/Caucasian (51.43%, n = 36), Black or African American (28.57%, n = 20), Asian (14.29%, n = 10), More than one race/ethnicity (2.86%, n = 2), American Indian or Alaskan Native (1.43%, n = 1), and 1.43% chose not to respond (n = 1). Study procedures were approved by the relevant University of Chicago Institutional Review Board and participants were compensated for their participation.

#### Stimuli

The food and nonfood stimuli were selected from the same source as Exp 1.

#### Procedure

Participants completed a source memory learning task designed to assess memory for both the identity of visual items and their spatial location (Zhao & Vogel, 2025). Each learning cycle consisted of a study phase followed by a test phase, and this study-test sequence was repeated five times. In each study phase, participants viewed 48 trials containing a mixture of food-related and neutral images. On each trial, an image was presented for 1000 ms in one of the screen quadrants, while a central fixation cross remained visible. Participants were instructed to memorize both the item and the quadrant in which it appeared. After the study phase, participants completed a source memory test phase, which also included 48 trials with food and neutral items.

Each test trial began with a 500 ms intertrial interval (ITI) showing a fixation cross. The image was then presented again, and participants responded without time limit to indicate which quadrant the image had appeared in during the preceding study phase. Thus, the task measured source memory accuracy for spatial context, rather than simple item recognition alone (see **Fig. 4**).

**Fig 4.**
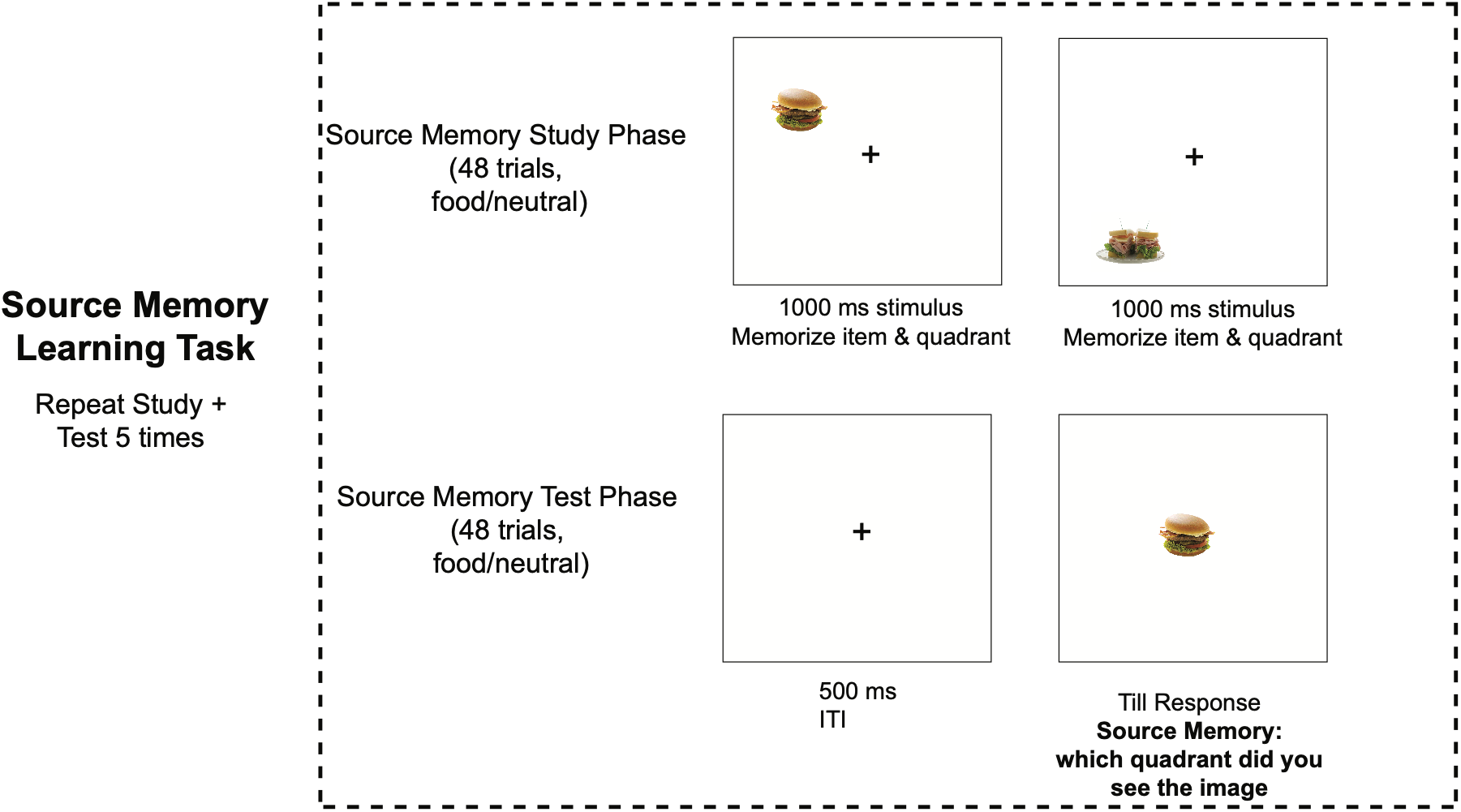
Schematic of the experimental paradigm used in Experiment 2. In each cycle, participants first completed a study phase in which 48 food and neutral images were presented for 1000 ms in one of four screen quadrants, and they were instructed to memorize both the item and its location. This was followed by a test phase of 48 trials, in which each trial began with a 500 ms ITI and participants indicated, with unlimited response time, which quadrant the image had previously appeared in. The study + test sequence was repeated five times.

**Table 2.** Mean, SD, Skewness and Kurtosis of Eating-Related Symptoms in Exp 2.

|  | Mean | SD | Skewness | Kurtosis |
|---|---|---|---|---|
| EPSI Restrictive Eating | 9.63 | 5.42 | 0.26 | -0.83 |
| EPSI Cognitive Restraint | 5.24 | 2.91 | 0.78 | 0.17 |
| NIAS Total Score | 15.01 | 10.38 | 0.27 | -0.74 |
| NIAS Picky Eating | 6.13 | 4.65 | 0.10 | -1.32 |
| NIAS Appetite | 4.94 | 4.14 | 0.54 | -0.46 |
| NIAS Fear of Negative Consequences | 3.94 | 4.04 | 0.75 | -0.55 |

### Results

#### Restrictive eating ratings were associated with lower recollection-based source memory accuracy for food relative to nonfood stimuli

We directly tested how eating preferences affect recollective memory. A mixed- effects model predicting source memory accuracy showed a significant main effect of time (b = 0.062, SE = 0.005, z = 13.39, p < .001), indicating that recollection-based source memory accuracy improved across repeated study-test cycles. There was also a significant negative main effect of restrictive eating propensity, as measured by the EPSI Restricting subscale (b = -0.060, SE = 0.024, z = -2.47, p = .014), such that higher restrictive eating was associated with lower source memory accuracy in the food condition, which served as the reference category.

Most importantly, this effect was qualified by a significant interaction between condition and EPSI Restricting (b = 0.022, SE = 0.009, z = 2.33, p = .020). This indicates that the negative association between restrictive eating and source memory accuracy was weaker in the nonfood condition than in the food condition. In other words, individuals higher in restrictive eating showed relatively poorer recollection of the spatial context of food stimuli, whereas this pattern was attenuated for nonfood stimuli.

To further test the interaction involving EPSI Restricting, we compared participants with median-split high versus low Restricting scores separately for food and nonfood stimuli. These follow-up t tests showed that individuals high in restrictive eating had significantly lower source memory accuracy for food stimuli than those low in restrictive eating (t(68) = -2.75, p = .008, Fig. 5 left), whereas the corresponding difference for nonfood stimuli was smaller and did not reach conventional significance (t(68) = -1.88, p = .065, **Fig. 5A**). This pattern lends support to the **Memory Suppression Hypothesis**, such that individuals high in restrictive eating appeared to show selectively reduced memory for food information, consistent with the idea that food cues may trigger avoidance or suppression processes that undermine recollection of episodic details across subjects.

**Fig 5.**
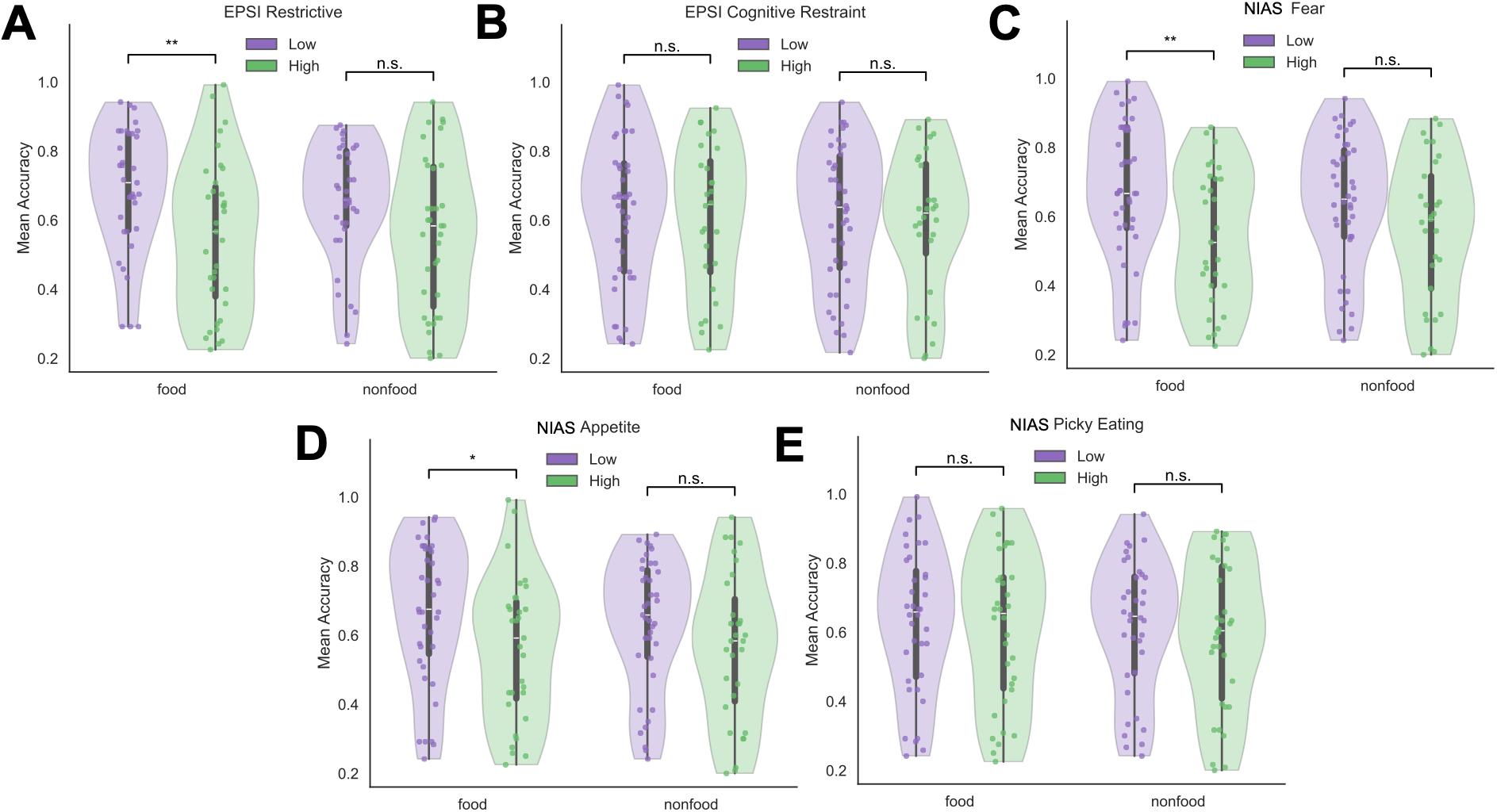
Restrictive eating behavior, fear of negative consequences, and lack of interest in eating are associated with reduced recollection of food relative to nonfood stimuli, whereas cognitive intent to control eating and picky eating are not. Higher EPSI Restring (A), higher NIAS Fear (C), and higher NIAS Appetite ratings (D) were selectively associated with reduced source memory accuracy for food images, but not for nonfood images. In contrast, other cognitive-control-related eating measures, including EPSI Cognitive Restraint (B) and NIAS Picky Eating (E), did not show differential recollection for food versus nonfood images between high and low groups.

Furthermore, there was a significant negative interaction between time and EPSI Restricting (b = -0.012, SE = 0.005, z = -2.48, p = .013), suggesting that individuals higher in restrictive eating showed a smaller improvement in source memory accuracy over time. However, the three-way interaction between time, condition, and EPSI Restricting was not significant (b = 0.006, SE = 0.007, z = 0.85, p = .397), indicating that the selective association between restrictive eating and poorer recollection for food versus nonfood stimuli did not significantly change across learning cycles.

Given support for the **Memory Suppression Hypothesis**, an important remaining question is whether the observed memory pattern is specifically tied to eating-related preferences or instead reflects a broader control mechanism. If this memory effect is specific to restrictive eating behaviors intended to control shape or weight, then cognitive intent to restrict food intake (EPSI Cognitive Restraint) would not be expected to show the same association with recollection. In contrast, if the effect reflects a broader process related to the affective salience of food, extending beyond shape-and weight-motivated restriction, then similar reductions in food recollection should also be observed among individuals with higher levels of cognitive restraint.

For EPSI Cognitive Restraint, the mixed-effects model predicting source memory accuracy showed a significant main effect of time (b = 0.062, SE = 0.005, z = 13.27, p < .001), indicating that source memory accuracy improved over time. However, there was no significant main effect of Cognitive Restraint (b = -0.024, SE = 0.026, z = -0.92, p = .359, **Fig. 5B**), suggesting that Cognitive Restraint was not associated with overall source memory accuracy in the food condition, which served as the reference category. Critically, the interaction between condition and Cognitive Restraint was not significant (b = -0.002, SE = 0.010, z = -0.24, p = .813), indicating that Cognitive Restraint was not differentially associated with source memory accuracy for food versus nonfood stimuli. The interaction between time and Cognitive Restraint was also not significant (b = 0.003, SE = 0.005, z = 0.62, p = .534), and the three-way interaction among time, condition, and Cognitive Restraint was not significant (b = 0.003, SE = 0.007, z = 0.41, p = .686). Overall, these results suggest that EPSI Cognitive Restraint was not a significant predictor of source memory accuracy, either directly or in interaction with condition or time.

For the NIAS total score, the mixed-effects model again showed a significant main effect of time (b = 0.062, SE = 0.005, z = 13.29, p < .001), indicating that source memory accuracy improved over time. However, there was no significant main effect of NIAS total score (b = -0.032, SE = 0.022, z = -1.46, p = .144), suggesting that the overall severity of ARFID symptoms was not associated with source memory accuracy in the food condition. The interaction between condition and NIAS total score was also not significant (b = 0.013, SE = 0.008, z = 1.55, p = .120), indicating that overall ARFID symptom severity did not significantly moderate differences in source memory accuracy between food and nonfood stimuli. Likewise, the interaction between time and NIAS total score was not significant (b = 0.003, SE = 0.004, z = 0.67, p = .502), and the three-way interaction among time, condition, and NIAS total score was not significant (b = 0.000, SE = 0.006, z = 0.08, p = .937).

When examining whether distinct ARFID subscales were associated with recollective memory performance, different patterns emerged. Higher Fear was associated with poorer recollective memory for food stimuli compared to nonfood stimuli (interaction between Fear x stimulus category, b = 0.016, SE = 0.007, z = 2.25, p = .025), with the high-Fear group performing significantly worse than the low-Fear group (t(68) = −2.75, p = .008, **Fig. 5C**) when presented with food stimuli, whereas no significant group difference was observed for non-food stimuli (t(68) = −1.54, p = .129). Picky Eating was not associated with recollective memory performance, as no significant differences were observed between the high- and low-Picky Eating groups for either food (t(68) = −0.41, p = .685, **Fig. 5E**) or non-food stimuli (t(68) = −0.44, p = .660). In contrast, higher scores on the Appetite subscale, indicating lack of interest in eating, were associated with worse recollective memory for food stimuli, with the high-Appetite group performing significantly worse than the low-Appetite group (t(68) = −2.08, p = .041, **Fig. 5D**) in this condition; however, the interaction between Appetite group and stimulus category was not significant (b = 0.015, SE = 0.009, z = 1.61, p = .107). No significant difference was observed for non-food stimuli (t(68) = −1.31, p = .195).

Overall, these findings suggest that ARFID-related fear of negative consequences and lack of interest in eating are associated with reduced recollective memory for food stimuli, an effect that was not observed for picky eating. Consistent with the EPSI Restricting subscale, these findings suggest that restrictive eating behavior, potentially driven by fear of negative consequences associated with eating and the lack of interest in eating, may be *selectively associated with a reduced recollection for food stimuli* rather than reflecting a broader cognitive control or avoidance mechanism.

## Discussion

This study examined whether stable individual differences in the personal significance of food influence long-term memory. Although affective state has been shown to enhance episodic memory, particularly recollection, this literature has largely relied on within-person manipulations of emotional valence. We instead asked whether differences in how individuals relate to food would predict memory for food relative to nonfood stimuli. Across two experiments, the results revealed a dissociation between familiarity and recollection. In Experiment 1, eating-related individual differences were largely unrelated to familiarity-based recognition memory. In Experiment 2, however, restrictive eating was associated with poorer recollection of food-related contextual information, and this effect was particularly evident among individuals reporting greater fear of negative consequences associated with eating. On the other hand, picky eating and cognitive restraint showed no comparable association with recollection for food. Together, these findings suggest that the mnemonic consequences of food-related affective significance are not uniform across memory processes. Rather, eating-related concerns appear to selectively influence recollection, with restrictive and fear-based responses to food associated with reduced access to episodic detail.

### Familiarity and Recollection Are Differentially Sensitive to Individual Differences in Eating-Related Behavior

The absence of a reliable relationship between eating-related individual differences and recognition memory in Experiment 1 provides an important constraint on accounts of affective influences on memory. Despite substantial individual variability in restrictive eating behaviors, cognitive restraint, and ARFID symptoms, none of these characteristics predicted familiarity-based recognition of food relative to nonfood stimuli. This pattern is consistent with the **Memory Independence Hypothesis** at the level of familiarity and suggests that differences in the personal affective significance of food do not necessarily alter the general familiarity of a food item.

This finding is notable given that the same individual differences were associated with differences in affective responses to food. Higher levels of restrictive eating behavior and ARFID symptoms were associated with relatively less positive evaluations of food than nonfood stimuli. Thus, eating-related differences were evident at the level of affective evaluation but did not translate into corresponding differences in familiarity-based memory. One possibility is that familiarity is relatively insensitive to the motivational or regulatory consequences of food-related affective significance.

Recognition can be supported by a relatively nonspecific sense of prior occurrence, whereas recollection requires the retrieval of richer episodic representations (Yonelinas, 1999, 2001). If restrictive eating involves the regulation of attention to, or engagement with, food-related information, such regulation may have little consequence for whether an item subsequently feels familiar while nevertheless compromising the retrieval of its contextual details.

This interpretation is consistent with the broader distinction between familiarity and recollection. Emotional manipulations often produce their strongest effects on recollection (Ochsner, 2000), suggesting that affective significance can influence the retrieval of detailed episodic representations without necessarily changing familiarity. The present findings extend this distinction by showing that the direction of the effect may depend on the nature of affective significance. Rather than producing a generalized enhancement of memory, food-related affective differences were associated with selective reductions in recollection.

An important consideration is whether these recollection deficits are already evident during the working memory encoding of food items. In contrast to Fukuda and Vogel (2019), however, our data revealed no set-size-dependent effect at the initial encoding stage. One possible explanation is that the food and nonfood stimuli used in the present study (i.e., animals) may have been more perceptually or semantically similar to one another than the common-object images used in the previous study (Brady et al., 2008). Consequently, although performance in our task was likely still constrained by working memory capacity during encoding, the similarity between stimuli may have introduced additional errors during the retrieval or comparison stage (Awh et al., 2007). Thus, the absence of a clear set-size effect may reflect errors arising during memory retrieval and comparison rather than a simple reduction in the number of items successfully encoded as set size increased. Furthermore, given the absence of significant interactions between category and the NIAS subscales, the present findings do not provide evidence that eating-related attitudes and behavior influenced the representation or familiarity of food versus nonfood objects in terms of working memory capacity or precision. This null effect may, however, be related to the categorical nature of the food stimuli used in the present study. Future research could examine food stimuli along a continuous perceptual dimension, such as color or shape (Zhang & Luck, 2008), to provide a more sensitive measure of changes in representational precision and to determine whether these changes interact with individual differences in eating behavior.

### Restrictive Eating Selectively Suppresses Recollection for Food

Experiment 2 provided evidence for the **Memory Suppression Hypothesis** with recollective memory. Higher levels of restrictive eating behavior, as measured by the EPSI Restricting subscale were associated with poorer source memory for food stimuli, and this relationship was weaker for nonfood stimuli. Individuals higher in restrictive eating therefore showed a relative deficit in remembering the spatial context in which food had been encountered. Importantly, this effect was observed in a task requiring memory for contextual information rather than simple item familiarity, providing a direct test of whether restrictive eating is associated with reduced episodic detail. Food may be highly significant to individuals with restrictive eating behaviors, but that significance may not be uniformly beneficial for memory. When significance is accompanied by avoidance, conflict, or regulatory concerns, the motivational relevance of food may promote processes that ultimately reduce the accessibility of food-related recollective information.

Our findings are consistent with within-subject studies suggesting that people may be less accurate in monitoring the source of negatively valenced information. For example, participants showed poorer source memory for negative than neutral items (Cook et al., 2007). One explanation is that heightened attention to emotional material may reduce the binding of contextual details, while leaving item memory relatively intact. This may also explain why our findings generalized across individuals: individuals who experience more negative emotion toward food may attend more strongly to food- related information, potentially reducing contextual binding and selectively impairing source memory for food. At the neural level, one study using event-related potential (Mao et al., 2015) found reduced neural signatures of recollection for negative items using an emotional source-memory paradigm. Our findings are also broadly consistent with within-subject evidence that emotional stimuli primarily affect recollection rather than familiarity (Kensinger & Corkin, 2003). However, our results suggest that the effect of negative emotion on recollection may depend on individual differences in emotional experience. Future work should therefore consider personal relevance, rather than universal valence classifications, when examining emotion and memory.

The learning results provide additional evidence of how restrictive eating may or may not influence the accumulation of episodic memory across repeated learning. Source memory improved substantially across successive study-test cycles, indicating that repeated exposure strengthened contextual memory. However, the EPSI Restricting × Stimulus Category × Repetition interaction was not significant, suggesting that restrictive eating did not alter the rate at which source memory improved across repetitions. Instead, the present findings are more consistent with an effect on the initial level of the memory trace rather than on the slope of learning. In other words, food-related deficits may reflect differences in the formation or accessibility of recollective representations at encoding or retrieval, rather than differences in the benefits derived from repeated retrieval practice or subsequent memory consolidation. Although these results do not support any strong conclusions about the specific stage at which restrictive eating exerts its influence, they suggest that the impairment is unlikely to reflect a general failure to accumulate memory across repeated learning.

### Memory Suppression for Food-Specific Recollection Is Not Simply a Consequence of General Cognitive Restraint

An important question is whether the relationship between restrictive eating and food memory reflects a broad cognitive-control mechanism. If so, measures of general eating-related restraint should be expected to show similar effects. The absence of such an effect for EPSI Cognitive Restraint argues against this interpretation. Cognitive Restraint was unrelated to source memory and did not interact with stimulus category or learning time. Thus, the memory effect observed for restrictive eating behavior, as measured by the EPSI Restricting subscale, does not appear to follow simply from the desire to exert deliberate cognitive control over eating.

This distinction is theoretically important because EPSI Restricting and EPSI Cognitive Restraint, although related, capture different aspects of eating pathology. The former reflects restrictive eating behavior, whereas the latter reflects the cognitive intention to monitor or control food intake. Our findings show that motivational influences can be experimentally distinguished from general positive affect and that these factors can have dissociable effects on cognitive control (i.e., Chiew & Braver, 2014). From this perspective, our findings suggest that merely endorsing a cognitive intention to restrict food intake may not be sufficient to alter the representation of food stimuli in working memory. Instead, food-related memory may be more strongly influenced when restriction is accompanied by fear of negative consequences from eating (e.g., weight gain, choking), negative affect, or personally meaningful food preferences. This possibility is consistent with evidence that individual differences in food motivation and eating behavior can modulate responses to food cues (Castellanos et al., 2009; Werthmann et al., 2011). Thus, our findings favor a more nuanced account that simply exerting higher cognitive control may not be sufficient to produce a food- specific memory effect. Instead, the effect may instead emerge when cognitive restraint is translated into actual restriction, fear, or altered affective responses toward food.

A primary limitation of the present study is that the sample comprised individuals across the spectrum from normative to preclinical levels of eating pathology, rather than a clinically diagnosed population. Accordingly, the psychometric properties and behavioral patterns identified in this study may not fully generalize to individuals with more severe or clinically significant presentations. Future research should seek to replicate and extend these findings in targeted clinical populations, including individuals formally diagnosed with Anorexia Nervosa, Avoidant/Restrictive Food Intake Disorder (ARFID), and other eating disorders.

## Conclusion

In conclusion, the present study demonstrates that individual differences in the personal significance of food may influence food-specific long-term memory. Across two experiments, eating-related preferences were not associated with familiarity-based recognition, but restrictive eating behavior was linked to poorer recollection of contextual details. This relationship was especially pronounced among individuals who reported greater fear of negative consequences related to eating. In contrast, picky eating and cognitive restraint were not associated with reduced recollection. These findings suggest that food-related affective concerns may have distinct effects on different components of memory, particularly by disrupting the encoding or retrieval of detailed episodic information rather than general recognition. Overall, the results highlight the importance of considering individual differences in food-related attitudes and behaviors when examining the relationship between affect and memory.

## Acknowledgements

This research was supported by the National Institutes of Mental Health grant R01MH132792.

